# Convergent evolution as an indicator for selection during acute HIV-1 infection

**DOI:** 10.1101/168260

**Authors:** Frederic Bertels, Karin J. Metzner, Roland Regoes

**Author notes:** **Correspondence:** Max Planck Institute for Evolutionary Biology, August-Thienemann-Straße 2, 24306 Plön, Germany. **Cite as:** Bertels F, Metzner KJ, Regoes RR. Convergent evolution as an indicator for selection during acute HIV-1 infection. bioRxiv 168260, ver. 4 peer-reviewed and recommended by PCI Evolutionary Biology (2018). DOI: 10.1101/168260.

## Abstract

Convergent evolution describes the process of different populations acquiring similar phenotypes or genotypes. Complex organisms with large genomes only rarely and only under very strong selection converge to the same genotype. In contrast, independent virus populations with very small genomes often acquire identical mutations. Here we test the hypothesis of whether convergence in early HIV-1 infection is common enough to serve as an indicator for selection. To this end, we measure the number of convergent mutations in a well-studied dataset of full-length HIV-1 *env* genes sampled from HIV-1 infected individuals during early infection. We compare this data to a neutral model and find an excess of convergent mutations. Convergent mutations are not evenly distributed across the env gene, but more likely to occur in gp41, which suggests that convergent mutations provide a selective advantage and hence are positively selected. In contrast, mutations that are only found in an HIV-1 population of a single individual are significantly affected by purifying selection. Our analysis suggests that comparisons between convergent and private mutations with neutral models allow us to identify positive and negative selection in small viral genomes. Our results also show that selection significantly shapes HIV-1 populations even before the onset of the adaptive immune system.

## Introduction

Convergent evolution is ubiquitous in nature. Phenotypic convergence is the repeated and independent evolution of a particular phenotype. Textbook examples are the evolution of flight, which occurred in bats, birds and insects independently, or the evolution of the eye, which arose many times independently across the tree of life (Serb and Eernisse 2008). We can also observe convergent evolution at the level of the genotype, which usually occurs when organisms are under extremely high selection pressure. Well known examples are the evolution of antibiotic resistance through the acquisition of mutations in the same gene or even at the same position in that gene (Farhat et al. 2013; Lindsey et al. 2013) or the evolution of cancer, which usually involves a driver mutation in a typical cancer or cancer suppressor gene (Miki et al. 1994). Convergent evolution in viruses is particularly common due to the small and functionally constrained viral genomes (Bull et al. 1997; Crandall et al. 1999; Wichman et al. 2000; Xue et al. 2017).

To study convergent evolution in viruses we focus on one of the best studied human viruses: Human immunodeficiency virus type 1 (HIV-1). HIV-1 is an RNA virus with a small genome and an extremely high mutation rate. The high mutation rate allows the virus to evade the human immune response and persist for years within the host (Coffin 1995; Koenig et al. 1995; Borrow et al. 1997; Goulder et al. 1997; Wei et al. 2003; Trkola et al. 2005; Liao et al. 2013). Over this time span HIV-1 evolves and acquires a large number of mutations (Shankarappa et al. 1999; Lemey et al. 2006; Keele et al. 2008; Poon et al. 2010).

During early infection of HIV-1 when immune escape does not seem to play a major role for HIV-1 evolution, HIV-1 evolution has been modeled as a neutral process (Lee et al. 2009; Giorgi et al. 2013). Similarly, diversification rates of the virus population have been shown to largely adhere to a molecular clock (Herbeck et al. 2011; Park et al. 2016). However, the adherence to a molecular clock does not mean that there is no selection during early HIV-1 infection. It is more likely that the fast accumulation of neutral mutations obscures selective footprints. Selection is more visible once viral phenotypes are taken into account. Typical phenotypes considered include the set-point viral load, which designates the average level of the virus during the chronic disease stage, or disease progression. For example, high evolutionary rates have been shown to correlate with fast disease progression and strong selection has been shown to correlate with slower disease progression (Boutwell et al. 2010; Garcia-Knight et al. 2016). Hence the data suggests that strong immune selection constrains viral diversification and hence leads to low evolutionary rates, which in turn leads to lower set point viral loads and therefore lower levels of disease progression.

Selection can be measured in various different ways: (1) Probably the most common way of determining selected nucleotide sites is by comparing the evolutionary rates of non-synonymous nucleotide sites to those of synonymous sites (dN/dS) on a phylogenetic tree (Kosakovsky Pond et al. 2008; Wood et al. 2009; Boutwell et al. 2010; Yoshida et al. 2011). (2) If population sequence data that spans multiple time points is available then it is possible to identify positively selected sites by assessing the change in mutant frequency over time (Henn et al. 2012; Foll et al. 2014). (3) More recently it has been demonstrated that it is possible to determine nucleotide sites under selection by analyzing the distribution of those sites across a gene (Zhang and Townsend 2009; Zhao et al. 2017).

Here we will focus on yet another way of determining nucleotide sites affected by selection: measuring the frequency of convergent mutations across different HIV-1 populations from different hosts (i.e. we define an HIV-1 population as all viruses from the same infected individual). Convergent mutations are mutations that occur in independent HIV-1 populations in parallel. More specifically convergence requires that two populations that share the same nucleotide at a specific position in the genome acquire the same mutation. However, due to the small HIV-1 genome it is possible that convergent mutations are the result of chance and not selection. Appropriate null models are necessary to distinguish adaptive from neutral mutations (Stayton 2015). Once we have made the distinction between selected and accidental convergence we reserve the term convergent mutation for mutations that occur in parallel in different populations.

By comparing mutations from HIV-1 populations to a random null model, we identify convergent mutations that are positively selected for. To this end we reanalyze a dataset of full length *env* genes sampled from infected individuals during early infection (Keele et al. 2008; Li et al. 2010). We find that some mutations are overrepresented in the Keele and Li datasets compared to a neutral model. These convergent mutations are significantly skewed towards gp41. This biased distribution of mutations indicates that convergent mutations are positively selected. In contrast, the biased distribution of private mutations (mutations that are found in a single HIV-1 population only) towards high diversity sites in the *env* gene suggests that private mutations are strongly affected by purifying selection. Hence our results show that HIV-1 is under a range of selection pressures immediately after infection of a new host even before the onset of the adaptive immune system.

## Results

We reanalyzed two datasets from previous studies on the evolution of HIV-1 during early infection of a single founder virus. The samples are estimated to have been taken less than 50 days after infection (Keele et al. 2008; Li et al. 2010). In total there are 95 *env* sequence alignments each from a single HIV-1 infected individual.

### Convergent mutations are unlikely under a neutral model of evolution

We observed convergent evolution for a large number of the 1059 mutations identified in the Keele and Li datasets (**Figure 1**, **Supplementary Data 1**, **Supplementary Data 2**). We compared the mutations observed in the Keele and Li data to 1000 random distributions of the same number and identical substitution rates as observed in the data (**Supplementary Figure 1**, **Figure 1**). When we redistributed mutations we distributed the same number of mutation we observed in each infected individual across the genome of this individuals HIV-1 consensus sequence. We also maintained transition and transversion rates of the original mutations. In a comparison with the neutral model we found that mutations occurring in three or more populations in parallel are overrepresented in the Keele and Li dataset. Mutations occurring in more than five HIV-1 populations in parallel are even more inconsistent with the neutral model. The number of convergent mutations declines linearly on a log scale for the neutral model. We do not see a linear decline for the Keele and Li data. Instead the decline levels off for highly convergent mutations (**Figure 1**). For example, there is one mutation that occurs in seven individuals in parallel, the likelihood to observe this in a neutral model is about 0.001. For lower levels of convergence the deviation between neutral model and observation is still significant. For example, in our simulations we observe on average one mutation that occurs in four HIV populations in parallel. In contrast, there are four such mutations in the Keele and Li data (**Figure 1**).

**Figure 1.**
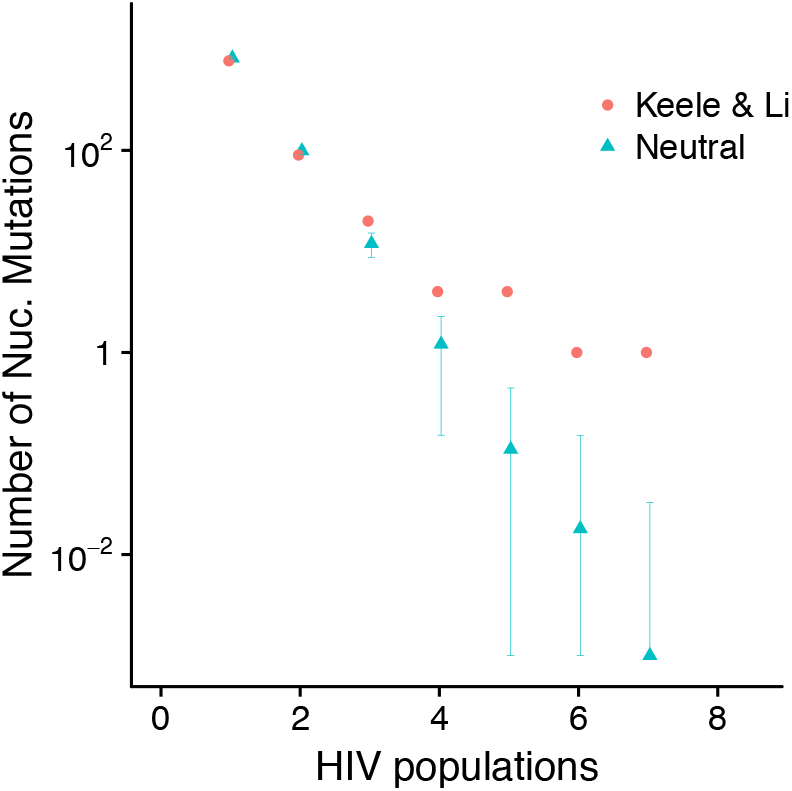
Convergent mutations are unusually frequent during early infection of HIV-1. Comparison between nucleotide mutations observed in one or more HIV-1 populations. Each HIV-1 population is isolated from a different infected individual. The Keele and Li data is compared to a neutral model of the same number of mutations as observed for each HIV-1 population using the same substitution rates as measured from the Keele and Li data. Data generated under the neutral model show the mean and standard deviations of 1000 simulations.

In total we determined 10 mutations that occur in four or more HIV-1 populations for which we expect to see a total of only one mutation under a neutral model of evolution (**Table 1**). Among these mutations there is one synonymous convergent mutation, which occurs in five populations in parallel. Mutations that occur in three populations in parallel are also overrepresented. We expect about 12 such mutations under our neutral model but observe 20 in the Keele and Li data. In 5 out of 1000 neutral models we observed 20 or more mutations occurring in parallel. Hence, probably some of the 20 mutations are positively selected. The exact identity and number of these mutations cannot be determined by considering parallelism.

**Table 1.**
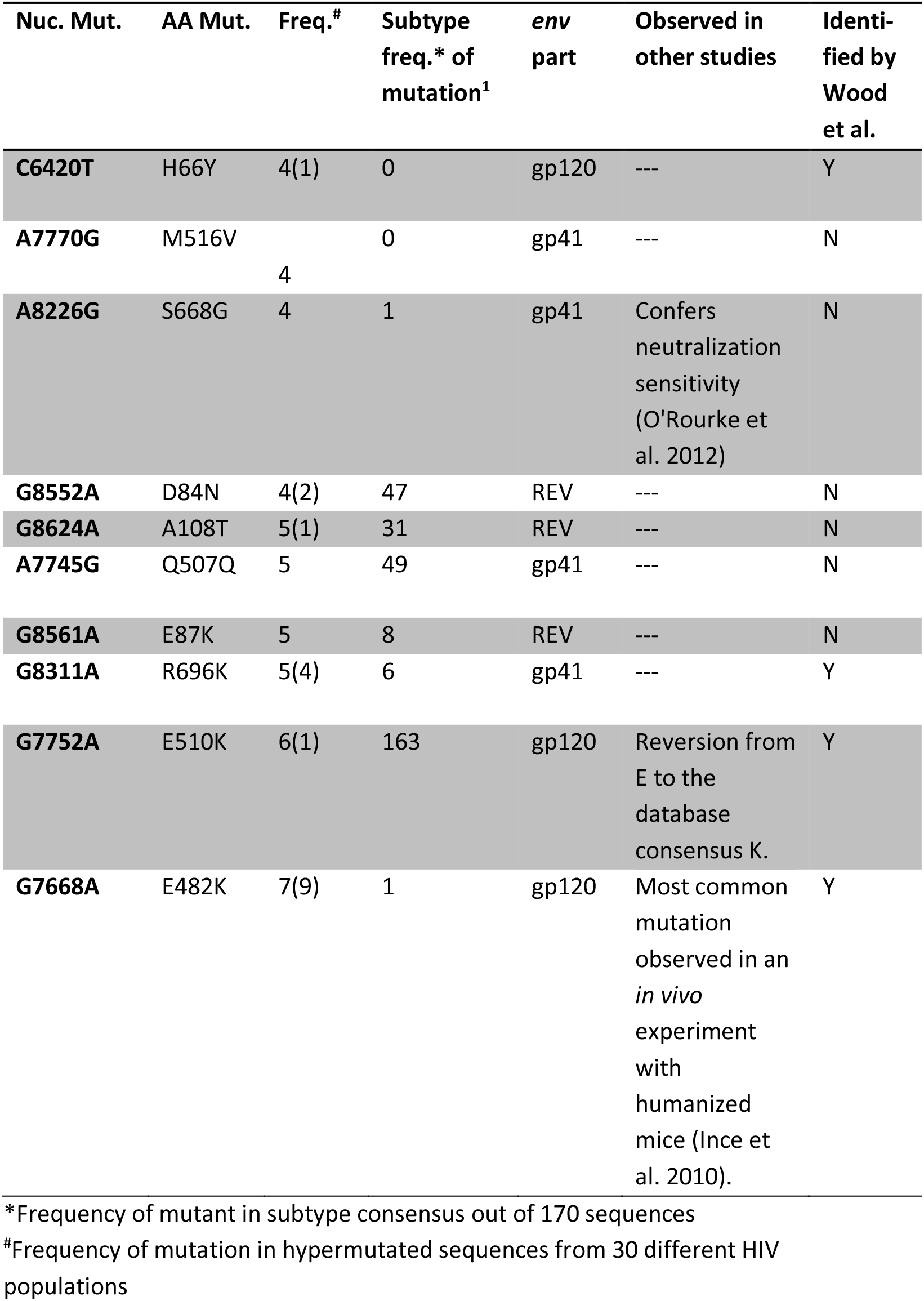
Convergent mutations occurring in four or more HIV-1 populations in parallel.

Despite the exclusion of hypermutated sequences the majority of the convergent mutations are G to A mutations (observing 6 out of 10 G to A mutations occurs in only 808 out of 10,000 random trials when we randomly draw mutations out of all observed mutations in the Keele and Li data, p=0.0808). Even though G to A mutations are not significantly overrepresented the number of G to A mutations is high and it is possible that residual or low APOBEC activity has caused some of these mutations. To test this hypothesis we analyzed the mutations that occur in all hypermutated sequences (sequences that contain four or more mutations) contained in the full dataset that we have excluded from the previous analysis. Only 30 HIV populations contain sequences with four or more mutations. The mutations in these sequences are predominantly G to A mutations (65% compared to 36% in the data lacking hypermutated sequences). Despite the low number of HIV populations that contain hypermutated sequences, mutations are shared in this dataset. The most common mutation occurs in nine HIV populations and is also the most common mutation in the dataset lacking hypermutated sequences (G7668A). The next two most common mutations occur in four HIV populations in parallel, one of them also occurs in the dataset lacking hypermutated sequences (G8311A). In total, six of the 10 convergent mutations also occur in hypermutated sequences. Of these six mutations, three occur in the hypermutated dataset in two or more HIV populations. If we exclude those three mutations as these may still be the result of residual APOBEC activity then there are seven mutations that are likely the result of positive selection. Among these seven mutations three are G to A mutations. Observing three or more G to A mutations is a very likely outcome of a sampling experiment when using the observed substitution rates (6551 out of 10,000 trials, p=0.6551).

### Convergent mutations occur predominantly in gp41

Convergent mutations are also differently distributed across the env gene compared to mutations that only occur in a single HIV-1 population (private mutations). Convergent mutations are more likely to be located at the end of the env gene (Figure 2). Private mutations are relatively evenly distributed across the entire env gene. The mean of the positions of private mutations is 439, which is very close to half the length of the env gene (428). However, the mean increases for mutations that occur in more than one HIV-1 population.

**Figure 2.**
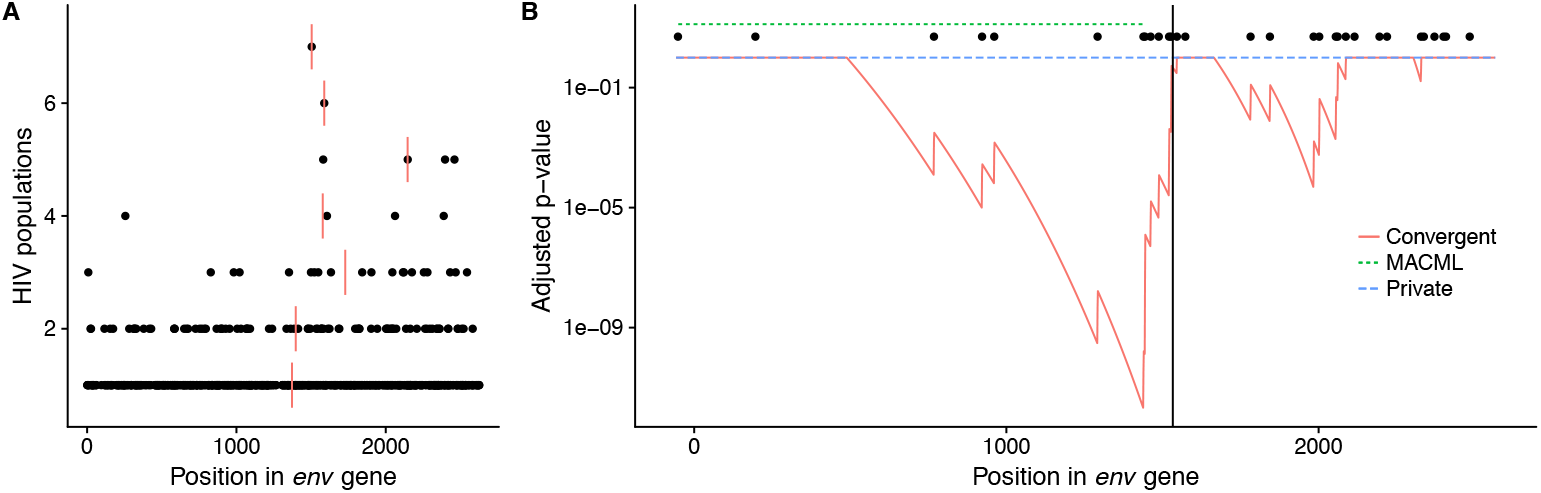
Convergent mutations are predominantly located in gp41. (A) Shows the positions of mutations across the env gene on the x-axis and the number of HIV-1 populations the mutations were observed in on the y-axis. Red bars indicate the mean of all positions of the mutations that occur in the same number of HIV-1 populations. (B) Shows whether there is a significant difference (adjusted p-value on y-axis) in the distribution of mutations before a certain position in the env gene compared to after this position. There is no significant signal for the distribution of private mutations (blue line). The red line shows the adjusted p-values for the distribution of mutations that occur in at least three HIV populations in parallel. The lowest p-value is found at nucleotide position 1438 in the env gene (position 7663 in HXB2, 479 in the env protein). For convergent mutations, a low mutation density region was identified by MACML (Zhang and Townsend 2009) and is indicated in green. The end of this region is also the minimum of the adjusted mutation distribution p-value from above. The black vertical line indicates the start of the Gp41 protein, a fusion protein that is part of the env gene. Black dots above the blue line indicate the positions of mutations that occur in three or more HIV-1 populations.

When dividing the *env* gene at every possible position and comparing the number of mutations observed before and after this position then we find, again, that there is not a single position that shows a significant difference between the distribution of mutations for the 5’ and 3’ end of the gene after adjusting for multiple testing (**Figure 2B**). However, mutations that occur in three or more HIV populations show very significant differences in their distribution. We found the largest difference at position 479 (adj. p-value 1e-12), separating the 5’ part of the *env* gene with few mutations from the 3’ part with many mutations. The 3’ part of the *env* gene encodes for Gp41, a protein responsible for forming the six-helix bundle during cell fusion with the host cell (Chan et al. 1997; Skehel and Wiley 1998; Melikyan et al. 2000).

### Private mutations are more likely to cause synonymous changes

The proportion of synonymous mutations in the Keele and Li data differ significantly from the predictions of a neutral model for private mutations (Figure 3). For private mutations we observe more synonymous mutations in the Keele and Li data than expected under a neutral model. This phenomenon can probably be explained by purifying selection. Purifying selection has probably led to the disappearance of non-synonymous mutations due to lethal or highly deleterious effects. The disappearance of lethal and highly deleterious mutations then leads to an overrepresentation of synonymous mutations in the data set. Generally synonymous mutations are more likely to have less deleterious effects because they do not change the amino acid sequence of proteins.

**Figure 3.**
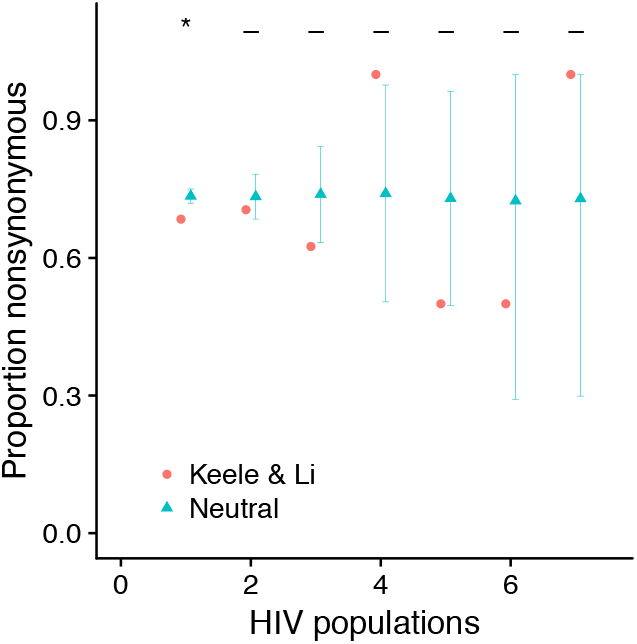
Private mutations show lower proportions of non-synonymous mutations than expected by chance. The proportion of synonymous mutations in the Keele and Li data compared to a neutral model (randomly distributed mutations, 1000 simulations). For each randomization the number of mutations in each convergence category (x-axis) was kept constant. A means there is no significant difference between the neutral model and the Keele and Li data, whereas a “*” means that the proportion of non-synonymous mutations in the Keele and Li data was lower in at least 950 out of 1000 simulations. Mutations in overlapping reading frames are excluded. The error bars show standard deviations.

### Private mutations are found significantly more frequently in regions of high diversity

Private mutations are mutations that we only see in a single HIV population in our dataset. These mutations are not simply a sample of all mutations that occur during the replication of HIV. Rather, because viral genomes replicate for more than one generation, the appearance of a mutation is the result of mutation and selection (see measuring mutation rates literature (Mansky 1996)). Selection will act because mutations that, for example, introduce stop codons in the middle of an essential gene will be lost in the next generation. Those kinds of mutations we expect to be underrepresented in our set of 770 env mutations. The distribution of those 770 mutations across the env gene does not seem particularly biased, as clustering approaches have shown (Figure 2).

Other than the location of mutations across the gene, we can also measure nucleotide diversity at each site in the env gene. We measure nucleotide diversity in an alignment of all 95 different consensus sequences. For these measures we do not take the mutations that we have observed in the Keele and Li data into account. For comparison to a neutral model, we randomly distribute mutations across the entire *env* gene and measure the mean diversity across all 770 positions. Interestingly the mean diversity of randomly distributed mutations in 1000 independent simulations was always lower than the mean diversity at the positions, at which the Keele and Li mutations occurred (**Figure 4**). This means that mutations do not occur in low diversity regions in the Keele and Li data. We propose that this pattern is caused by purifying selection, i.e. when mutations occur at low diversity sites they cause strongly deleterious or lethal phenotypes that will not leave offspring in the viral population and hence will be underrepresented in the Keele and Li dataset.

**Figure 4.**
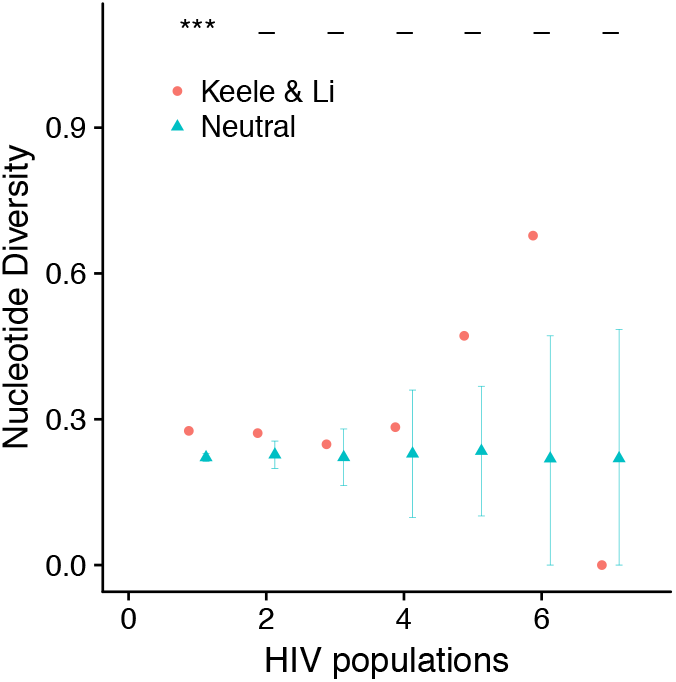
Private mutations occur in positions of high nucleotide diversity. The figure shows the nucleotide diversity (mean and standard deviation) at nucleotide positions, for which Keele and Li mutations were identified (orange) as well as the positions for 1000 individual random mutation distributions (turquois). The positions of the same number of mutations we observe in *N* HIV-1 populations was randomized in the neutral model. The diversity was determined at each position in the *env* consensus sequence alignment of all 95 HIV-1 populations (see Methods). A diversity of 1 means that all nucleotides observed at a certain position occur at equal frequencies. “***” indicates that among 1000 simulations there was not a single simulation that showed a mean diversity higher or equal to that observed in the Keele and Li data. “—” indicates that there was no significant difference between the diversity observed in the Keele and Li data and 1000 random distributions of mutations.

### Non-synonymous private mutations occur in highly diverse regions of the env gene

The significantly higher proportion of synonymous mutations does not cause the significantly higher nucleotide diversity for private mutations. We can understand the causal relationship by comparing the nucleotide diversity of synonymous and non-synonymous mutations and their effect sizes for randomly distributed mutations.

The nucleotide diversity of all private Keele and Li mutations is on average 0.26, the mean diversity of private mutations from a set of randomly distributed mutations is 0.22 (the maximum of all 1000 randomizations is 0.24, **Figure 5**). The nucleotide diversity at synonymous sites increases from a mean of 0.29 for randomly distributed mutations to 0.3 in the Keele and Li data. The mean diversity of non-synonymous sites increases from 0.19 in the randomization to 0.24 in the Keele and Li data.

**Figure 5.**
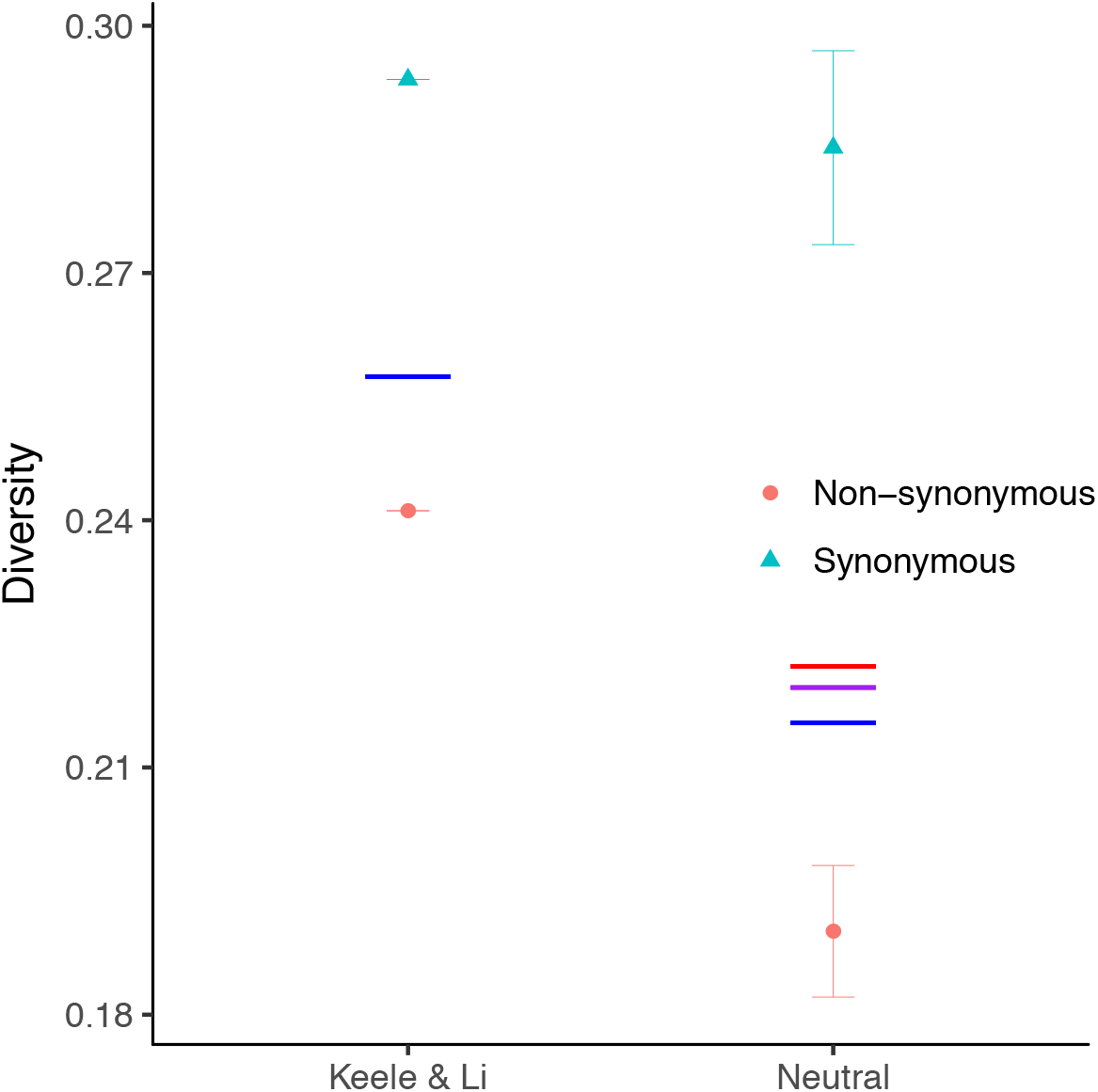
Nucleotide diversity of private non-synonymous and synonymous mutations in the Keele and Li data compared to our neutral model. The blue lines indicate the mean diversity of all private mutations. The purple line indicates the diversity of the neutral model if we adjust the proportion of synonymous mutations to that observed in the Keele and Li data. The red line shows the mean diversity of randomly distributed mutations if the proportion of synonymous mutations and the diversity of synonymous mutations is set to that observed in the Keele and Li data. Error bars show standard deviations.

Hence, non-synonymous mutations deviate much more from the neutral expectation than synonymous mutations, which are almost identical to the neutral expectation.

If we set the proportion of synonymous sites in one of our random simulations to the value observed in the Keele and Li data then the nucleotide diversity increases only slightly from 0.217 to 0.221 (purple line in **Figure 5**). If we instead increase the diversity of non-synonymous mutations to that observed in the Keele and Li data, then the mean diversity increases by 0.04, a ten-fold difference in effect size compared to increasing the proportion of synonymous mutations. This demonstrates that the increased proportion of synonymous mutations is not the main driver for the significantly higher nucleotide diversity of private mutations compared to our neutral simulation. Instead, the differences in diversity can be explained by non-synonymous mutations that are more likely to occur at highly diverse sites of the *env* gene (see also **Supplementary Figure 2**).

If purifying selection is the cause for the observed distribution of non-synonymous mutations then non-synonymous mutations do not preferentially occur in high diversity regions of the gene (although mutational hot spots could also cause part of the signal). Instead non-synonymous mutations in conserved regions are absent from the data because these mutations are likely to cause highly deleterious changes, which immediately disappear from the population. We do not see this bias in synonymous mutations because synonymous mutations are more likely to cause neutral or only slightly deleterious changes. Hence, it is more likely for us to observe the complete spectrum of synonymous mutations.

Therefore we can infer that non-synonymous mutations in low diversity regions have very strong negative effects on viral fitness. As one would expect these effects are much smaller for synonymous mutations where the difference between neutral model and Keele and Li data is much smaller (**Figure 5**). Nevertheless, there is a difference (although small) between the neutral model and Keele and Li data for synonymous mutations and we also identify one synonymous convergent mutation (**Table 1**) suggesting that synonymous mutations can have non-neutral effects on HIV-1 evolution.

## Discussion

Convergent evolution, here defined as acquiring identical mutations in independent evolving populations, can be a good indicator for selection when a large number of independent populations can be sampled. In our example, mutations that emerge more than three times in independent HIV populations are very unlikely to occur in parallel by chance. Furthermore convergent mutations occur preferentially in gp41, which is not the case for private mutations and hence supports the hypothesis that convergent mutations are positively selected.

Gp41 is the C-terminal part of Env and responsible for fusion with the host cell (Chan et al. 1997; Skehel and Wiley 1998; Melikyan et al. 2000). It is conceivable that increased fusion efficiency with the host cell provides significant fitness benefits even during exponential growth in the early stages of infection by HIV-1. An example for one of these mutations is E510K. This mutation is a reversion from a relatively rare amino acid to the database consensus (Foley et al. 2013; Davey et al. 2014). Reversions to ancestral states have been observed to continue for years after infection, consistent with our observation (Carlson et al. 2014). The most common mutation E482K (occurs in seven infected individuals) is not in gp41 but has also been observed in a previous experiment as the most common mutation in a humanized mouse experiment (Ince et al. 2010). However, we also find that E482K is even more common in hypermutated sequences (occurs in nine out of 30 HIV populations), which could mean that E482K is induced by APOBEC activity and may not be positively selected.

Wood et al. identified 36 *env* positions that are potentially under strong positive selection (Wood et al. 2009) mainly using methods based on dN/dS measurements. We identified a total of 10 mutations that occurred in four or more populations and among these seven that likely arose due to positive selection (**Table 1**). Only four mutations are shared between our list and the list published by Wood et al. We identify five novel putatively selected sites one is a synonymous site and hence could not have been identified with dN/dS. There are several reasons for the small overlap between the mutations identified by us and by Wood et al. First, because we wanted to exclude mutations caused by APOBEC as well as sequences that are unlikely to replicate due to a high mutational load, we excluded all sequences that contained three or more mutations compared to the consensus sequence. Second, Wood et al. have not included the Li dataset and hence only used 81 infected individuals instead of 95 in their analysis. Third, it is likely that some non-convergent mutations are also positively selected. These mutations may be beneficial only in a particular host and hence it would be impossible for our analysis to identify such mutations. Fourth, we have not included mutations that occur in three HIV-1 populations in parallel in our list, although about eight of those mutations are potentially the result of positive selection. Wood et al also identified one of the 20 mutations that occur in three populations in parallel (E322K) as under positive selection.

Although an excess of convergent mutations compared to a neutral model is a good indicator of selection, the strength of selection is difficult to infer from the number of convergent mutations. A mutation that occurs in a large number of individuals may be a mutation that is beneficial in a large number of different environments. Such a mutation may still have a smaller fitness effect than a mutation that increases the viral fitness only in a single environment but deleterious in the rest. Nevertheless, it is likely that the convergent mutations we identified have large fitness effects, as they need to have significantly increased in frequency since infection to be present in the Keele and Li dataset. It is possible to explicitly simulate the evolution of HIV-1 and to thus infer fitness effect distributions of convergent mutations (Bons et al. 2018).

One major advantage for identifying selected sites by analyzing convergent mutations over identification of selection in time course experiments is that linkage effects do not have to be considered. Mutations that rise in frequency in a population as the result of selection can sometimes also lead to sweeps of neutral or even detrimental mutations. However, the probability that an identical mutation hitchhikes multiple times independently with a beneficial mutation to fixation or high frequency should be similar to that of our neutral model. Hence hitchhiking mutations should not become convergent mutations.

Finally, our work also raises the question of why these mutations have not been present in the ancestral strain if they arose in parallel in up to 7% of all subjects. There may be multiple reasons for this phenomenon. First, it is possible that the observed mutations are only beneficial in a small proportion of all hosts. Second, it is possible that the convergent mutations we observed are beneficial during exponential growth but do not confer benefits after the immune response takes effect and hence will be lost during later stages of infection.

One could also wonder why there are only few mutations that occur in parallel in few hosts. This may be because mutations that are beneficial in most environments (human hosts) have probably already swept through the entire HIV-1 population and are now part of the consensus sequence. There may have been more convergent evolution early on during the HIV-1 epidemic when HIV diversity was low and the virus was still adapting to the human host.

## Methods

### Identification of mutations

The Keele and Li datasets consist of full-length env sequence alignments from a total of 102 and 30 infected individuals, respectively, containing on average 29 (minimum of 11 to a maximum of 63) full-length env sequences per infected individual amplified by single genome amplification (Keele et al. 2008; Li et al. 2010). The sequences are from viruses at different stages of early infection but each infected individual was sampled only once. In 78 of the 102 infected individuals from the Keele study and in 17 out of 30 infected individuals of the Li study the infection was likely caused by a single founder strain. We only analyzed those sequence alignments (a total of 95), because it becomes almost impossible to distinguish mutations from recombination events for the other cases (Keele et al. 2008).

We defined a mutation as a change from the consensus sequence of the virus population. The consensus sequence of each HIV-1 population is likely to be identical to the most recent common ancestor or even the sequence of the founder virus (Keele et al. 2008). Between infected individuals the consensus sequences differ. Hence to be able to compare mutations between individuals we also aligned all sequences to each other.

### Mutation comparison between HIV populations

To be able to compare mutations that occurred in virus populations from different infected individuals we aligned all of 95 alignments to each other as well as a reference sequence (HXB2). Alignments were performed using Clustal-Omega with standard settings (Sievers et al. 2011). Mutations that occur in virus populations of different infected individuals that changed the same consensus nucleotide to the same mutated nucleotide are considered identical.

### Neutral mutation distribution model

To assess whether the number of observed convergent mutations is indicative of selection we compared the Keele and Li data to a neutral model. We constructed the model in the following way:

1. We determined all possible mutations for each consensus sequence individually to generate a pool of mutations from which we can later randomly draw mutations.
2. To make sure that our results are not affected by mutation biases we amplified mutations according to their respective substitution rates measured from the Keele and Li dataset. First we counted the frequency of each of the twelve possible substitutions (A to C, A to G…). We normalized the lowest value in the substitution matrix to one and the remaining values in the matrix were scaled up accordingly. Each mutation from step one was amplified by the respective value in the substitution matrix.
3. This means for each of the 95 individual HIV-1 populations we end up with a set of possible mutations that are normalized by substitution frequency (e.g. G to A substitutions are more frequent than G to C mutations). From these normalized sets of mutations we randomly draw mutations without replacement (we cannot draw the same mutation twice). The number of mutations we draw is determined by how many mutations were originally observed in each of the 95 HIV-1 populations (identical mutations are only counted once!).
4. For the resulting dataset convergence analyses can be performed.
5. We repeated step 3) and 5) 1000 times to obtain statistically robust results.

### Randomly distributing mutations keeping the number of convergent mutations constant

To compare characteristics of convergent mutations to a neutral null model we randomized mutations for each convergence category (i.e. mutations occurring in different numbers of HIV-1 populations). In this randomization, we (1) maintain substitution rates; (2) maintain the origin of the mutation, i.e. mutations are randomly selected from the same HIV-1 population where they were observed in the Keele and Li dataset; and (3) only chose identical mutations if they were identical in the Keele and Li datasets.

### Measuring nucleotide diversity

Here we define nucleotide diversity as Shannon entropy at a certain position *i* for all consensus sequences.

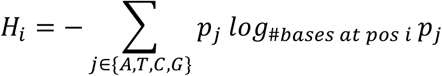

Where *p_j_* is the proportion of nucleotide *j* in all consensus sequences at reference position *i*.

The diversity at a particular position in the env gene was determined by calculating the Shannon entropy across all positions of a consensus sequence alignment. To normalize the diversity value to a range from 0 to 1, we determined the number of different bases present in all consensus sequences at each position and used this number as the base for the logarithm. Hence a diversity of 1 means that all observed nucleotides at a certain position occur in equal quantities. If only a single nucleotide occurred at a given position then the nucleotide diversity was set to 0.

#### Measuring the proportion of synonymous mutations

For each mutation we can determine whether it causes a synonymous or non-synonymous change. The proportion of synonymous mutations is the number of synonymous changes divided by the number of total changes. For **Figure 3** we excluded all positions from 8379 to 8653 in the HXB2 reference sequence because this region overlaps with the rev reading frame.

#### Measuring differences in the distribution of mutations across the env gene

We performed a chi-square test that tested whether the number of mutations before a given position is significantly different from the number of mutations after this position for the corresponding sequence lengths. This means we performed a total of 2623 chi-square tests. We then adjusted the p-values from these test for multiple tests with the Bonferroni Correction. All tests were used as implemented in R (Team).

Figures were created using cowplot and ggplot2 (Wickham 2016; Wilke 2016).

## Acknowledgements

This article has been peer-reviewed and recommended by *Peer Community in Evolutionary Biology* (https://dx.doi.org/10.24072/pci.evolbiol.100060).

## Supporting Information

**Supplementary Data 1. Position of all identified mutations in the *env* gene.** This file contains detailed information about the identity of the observed mutations. It provides the position in the HXB2 genome, the amino acid change they cause in the different genetic backgrounds and the number of HIV-1 subtypes (out of a total of 170) the mutations occurs in.

**Supplementary Data 2. Position of all identified mutations in the *rev* exon part of the *env* gene.** Same as Supplementary Data 1, except that only mutations and amino acid substitutions in the *rev* exon 2 are shown.

**Supplementary Figure 1.**
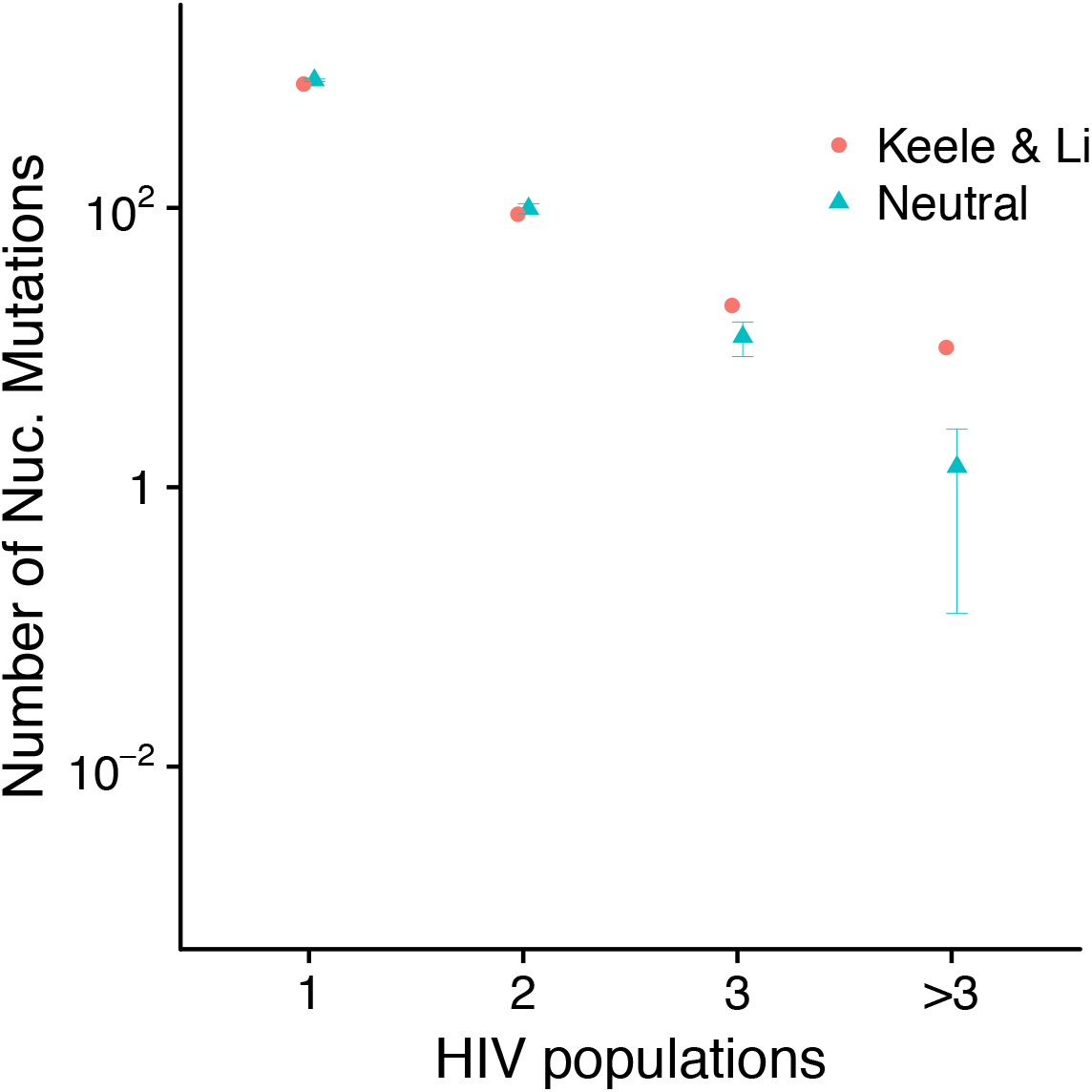
Convergent mutations are unusually frequent during early infection of HIV-1. Same as **Figure 1** except that mutations occurring in more than three HIV populations independently were grouped together.

**Supplementary Figure 2.**
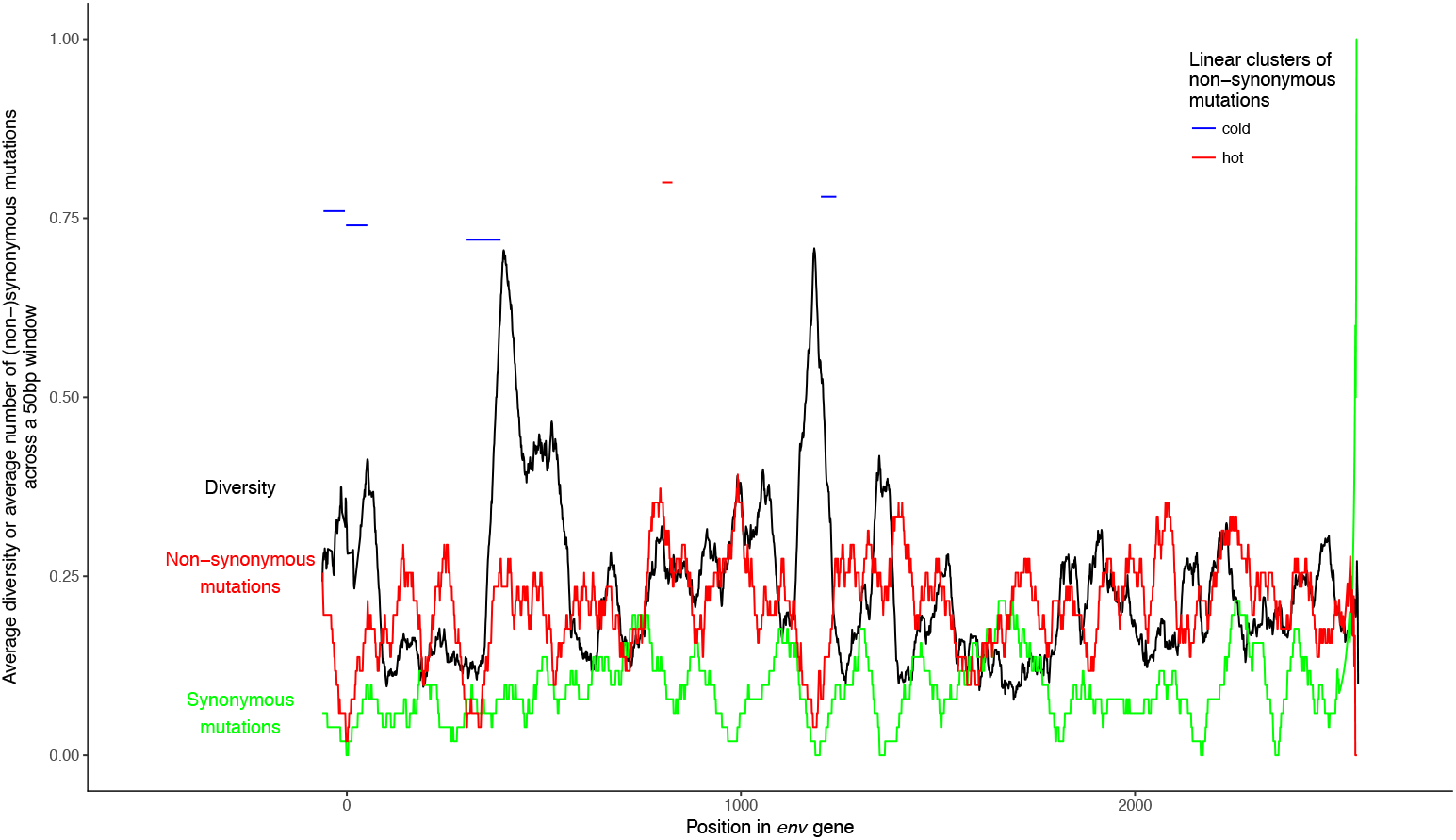
Correlation between the presence of nonsynonymous and synonymous mutations with diversity across the *env* gene. Linear clusters were computed with MACML (Zhang and Townsend 2009).

**Supplementary Program.** With this program one can redo the analyses and simulations of the manuscript.

